# Mechanical control of the insect extracellular matrix nanostructure

**DOI:** 10.1101/2024.08.20.608778

**Authors:** Yuki Itakura, Housei Wada, Sachi Inagaki, Shigeo Hayashi

**Affiliations:** Laboratory for Morphogenetic Signaling, RIKEN Center for Biosystems Dynamics Research, 2-2-3 Minatojima-minamimachi, Chuo-ku, Kobe, Hyogo, 650-0047, Japan; Department of Biology, Kobe University Graduate School of Science, 1-1 Rokkodai-cho, Nada-ku, Kobe, Hyogo, 657-8051, Japan

**Author notes:** **Correspondence to:** Shigeo Hayashi, RIKEN Center for Biosystems Dynamics Research, 2-2-3 Minatojima-minamimachi, Chuo-ku Kobe 650-0047, Japan.

## Abstract

Nanoscale modifications of apical extracellular matrix (ECM) have created a variety of functional surfaces with distinct physical properties, as exemplified by structural coloration and superhydrophobicity in animals and plants^1^. While the surface nanostructures of various organisms have inspired numerous biomimetic applications, the biological mechanisms underlying ECM patterning at the nanoscale remain largely unknown. Here, we investigated the morphogenesis of cuticular pores in *Drosophila* olfactory organs^2,3^. Hundreds of uniform-sized nanopores of ∼50 nm permit selective access of odorant molecules to olfactory neurons. We showed that matrices composed of the zona pellucida domain (ZPD) protein^4–6^ cover sensory organs in cell type-specific patterns and combinations. The ZPD proteins Dyl, Tyn, Mey, and Nyo form matrices with specific mixing and sorting properties, restricting cell growth and movement. The disruption of ZPD matrices leads to detachment of the envelope layer of the cuticle from the plasma membrane, and reduced numbers and irregularly sized nanopores. Our results suggest that compressive force from the ECM is essential for robust nanopore morphogenesis. This work reveals a previously unrecognized role for ZPD proteins as modular units that establish the mechanical environment required to modulate the nanoscale assembly of cuticle materials, opening a new biological context to these biomimetically important structures.

## Introduction

The outer surface of living organisms is coated with various forms of apical extracellular matrices (aECM), playing roles in protective functions and mechanical support. aECM is often modified to perform other specialized functions as well, such as selective light reflection, water repellency, molecular filtering in sensory organs, and surface ornamentation^1,2,7^. Structures of functional aECMs have been widely adopted in the biomimetic fabrication of inorganic materials for industrial uses^8,9^. A better understanding and control of nanoscale structures in biocompatible organic materials would thus greatly expand the potential for biomimetic research and applications^10,11^. However, development in this direction has been impeded by the lack of understanding of the fundamental biological mechanisms underlying ECM nanopatterning.

Cuticle, the aECM in insects, is a multi-layered structure comprising the envelope, epicuticle, and chitin-rich procuticle, which form in an outside-in order^12,13^. It has been used as the template for extensive evolutionary functionalization of diverse physical mechanisms, such as structural coloration and sensory functions. Wigglesworth (1948) suggested the crystallization of cuticular substances as a structural basis for iridescent coloration^12^. However, the role of self-organization in ECM nanopatterning has not been addressed.

Studying the nanopores, ∼50 nm diameter cuticular pores, that serve a molecular filtering function in the olfactory organ (Fig. 1a), we previously showed that the first sign of nanoscale structural modification appears at the earliest stage of cuticle formation, with the appearance of the wavy curved envelope (Fig. 1b, c), in which the bottom of the wave closest to the plasma membrane subsequently forms the nanopore^3^. *gore-tex/Osiris23 (gox/Osi23)*^3,14^ regulates this process by introducing the requisite “waviness” to the envelope. Folded/invaginated plasma membranes are frequently associated with prospective pore-forming regions^3^ (Fig. 1d), leading to the hypothesis that the wavy plasma membrane serves as the template for envelope curvature^3^. Since Gox/Osi23 protein is present in the endoplasmic reticulum (Inagaki et al., submitted), its influence on cuticle patterning is necessarily indirect. Further study of aECM materials and their mechanical properties is needed to gain a fuller mechanistic understanding of cuticle nanopatterning.

**Figure 1.**
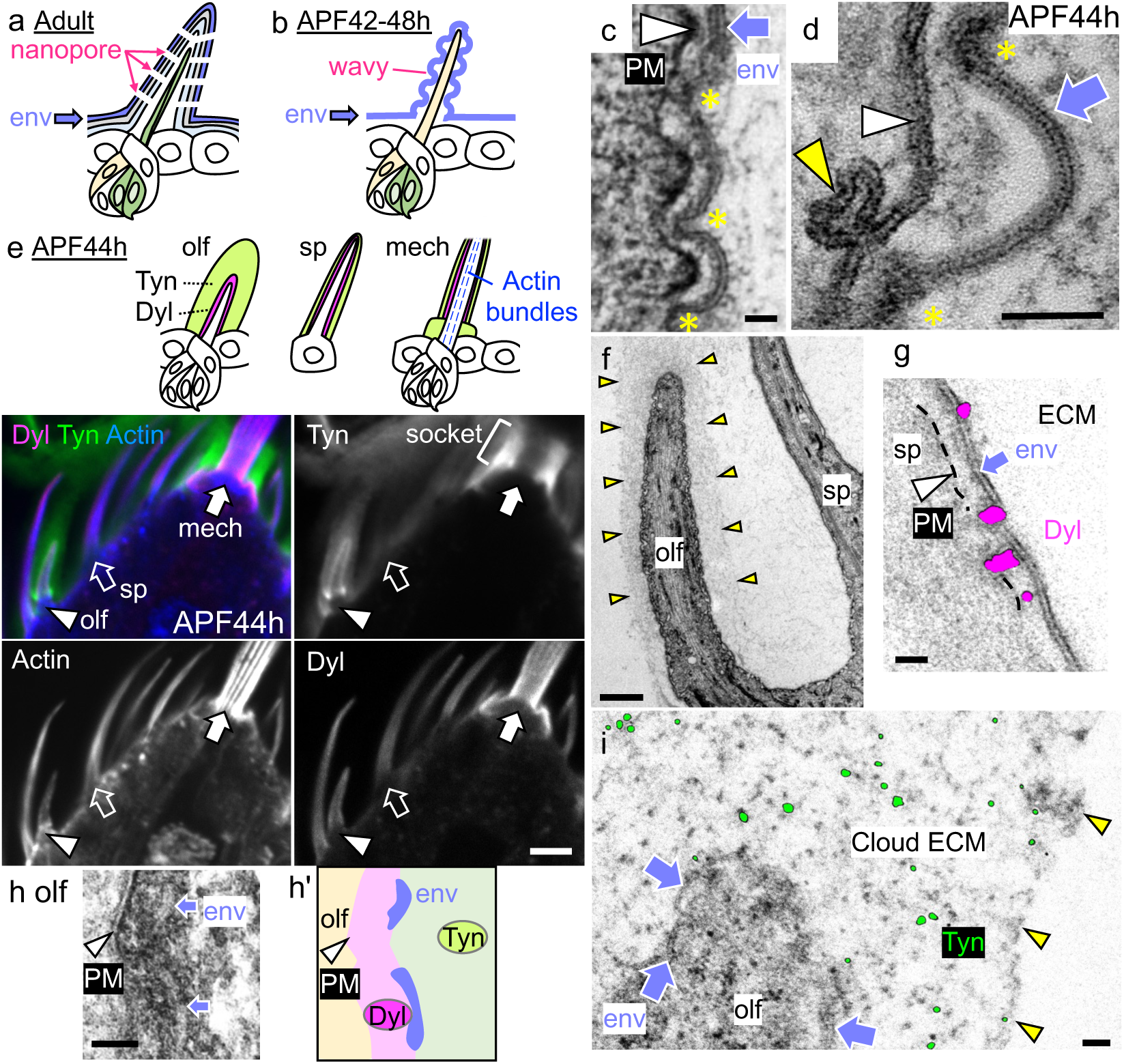
Layered ECM organization of the olfactory hair cell. **a,** A diagram of an adult olfactory bristle. The envelope (env) is the outermost layer of the cuticle. The nanopores allow odor molecules to reach the dendrites of sensory neurons (green). **b,** Olfactory hair cell at 42-48 h APF. The wavy env is formed above the plasma membrane of the hair cell (yellow). After cuticle formation, the hair cell retracts and is replaced by the dendrite. **c,** An electron-micrograph of a wavy env (arrow) over the wavy plasma membrane (PM, arrowhead). The innermost (yellow asterisks) and outermost parts of curved env and PM are placed in synchrony. **d,** The ladder-like env structure (arrow) with electron-dense innermost parts (yellow asterisk) and the lipid bilayer of PM (white arrowhead) are shown. The yellow arrowhead indicates a membrane invagination often observed close to the innermost part of the env. **e,** Top. The diagram of olfactory hairs (olf), spinules (sp), and mechanosensory bristles (mech) with localization patterns of Dyl, Tyn, and F-actin. Bottom. Fluorescence images of Dyl, Tyn, and F-actin staining. **f,** TEM image of olf and sp. The outer boundary of the cloudy material (cloud ECM) around the olf is indicated by yellow arrowheads. **g,** Immuno-EM visualization of Dyl (magenta) in sp. **h** and **h’** Ruthenium red staining of olf and the schematic diagram showing the position of plasma membrane, Dyl, envelope, and Tyn. **i,** Tyn signal (green) detected in the cloud ECM. Arrowheads indicate the outer boundary of cloud ECM. Scale bars: 5 μm for **e,** 1 μm for **f,** and 50 nm for the others.

ZPD proteins are conserved apical ECM components with the self-assembly property present in mammalian egg coats, anti-bacterial filaments in the urinary tract, tectorial membrane in the inner ear, and other biological processes^4,6,15^. In *Drosophila,* many ZPD proteins are expressed in epidermal tissues prior to cuticle formation in a cell type-specific manner and set up the cuticle shapes of tracheal tubules and epidermal denticles^16–19^. Here, we address the roles of ZPD proteins in cuticle nanopatterning, focusing on Dusky-like (Dyl), Trynity (Tyn), Morpheyus (Mey), and Nyobe (Nyo), all of which have been shown to play critical roles in cuticle formation^16,19–24^.

### Cell type-specific modular organization of ZPD proteins

The envelope layer emerges as a 20 nm-thick, ladder-like structure above the cell surface at ∼44 h after puparium formation (APF, Fig. 1b-d). We compared the ZPD protein distribution surrounding the olfactory hair cell of sensilla basiconica in the maxillary palp (olf) with that over the spinule (sp), a poreless epidermal protrusion, and the mechanosensory hair (mech), which produces a flat envelope and no nanopores.

Dyl and Tyn were visualized using protein-tagging and immunostaining techniques, and consistent patterns were observed (Fig. 1e, Extended Data Fig. 1a). Dyl formed a tight layer surrounding the surface of hair cells and epidermis, with enhanced signals in sp and mech (Fig. 1e). In contrast, Tyn was prominently enriched in the olf, forming thick layers covering the olf (Fig. 1e). Cloudy extracellular materials were also observed specifically on the olf under transmission electron microscopy (EM, Fig. 1f). We named this structure “Cloud ECM.”

Immuno-EM was performed to detect the localization of mVenus-tagged knock-in alleles of Tyn and Dyl (Extended Data Fig. 1b). Due to the high spatial resolution and the sporadic nature of signal detection by gold enhancement, the signal appeared as discrete spots. Dyl was detected mainly between the envelope and cell membrane of sp (Fig. 1g). Ruthenium Red (RR), which stains acidic mucopolysaccharides and glycosaminoglycans enriched in ECM, stained densely in the membrane-proximal zone and loosely the outside zone on the olf (Fig. 1h). At the border of the two zones, negatively stained convex structures around 50 nm above the PM, which are likely the envelope, were observed (Fig. 1h). Thus, RR stains Dyl region. Tyn signals were broadly distributed in the cloud ECM (Fig. 1i).

Other ZPD proteins, Nyo, Mey, and Neo, were detected on the olf, mech, and sp. Those proteins occupied distinct territories surrounding the olf (Extended Data Fig. 1c-e). aECM of the olf was subdivided into three layers. Layer I covers up to 125 nm above the cell surface, and is enriched with Dyl (Extended Data Fig. 1j). At 38 h APF, Tyn, Neo and Mey begin to cover layer I to form layer II. Tyn and a part of Mey expand further distally to form layer III. At 44 h APF, Nyo and Neo occupy layer II and III, respectively. The cloud ECM corresponds to layers II and III. The amounts of Tyn, Mey, and Nyo are low in the mech and sp. In summary, layer I (Dyl) covers all epidermis and its derivatives in 44 h APF. Olf is specifically covered by cloud ECM (layer II and III).

### ZPD proteins self-organize to build distinct ECMs

To uncover how ZPD proteins form ECM compartments on an olf hair cell, we expressed them in macrophage-derived *Drosophila* Schneider 2 (S2) cells expressing almost no ZPD transcript^25^. Separately expressed Tyn, Dyl, Nyo, and Mey were secreted and formed stable ECM structures. Those patterns were maintained when expressed in combinations (Fig. 2a, d–h, Extended Data Fig. 2). Neo was not secreted and was thus not analyzed further (Extended Data Fig. 2i). Tyn formed loosely distributed matrix, and Dyl formed thin shell-like layer covering the plasma membrane (Fig. 2a). Immunoprecipitation analysis of Tyn and Dyl, expressed in different S2 cells, revealed that the two proteins predominantly formed homo-oligomers, with some forming hetero-oligomers (Fig. 2b).

**Figure 2.**
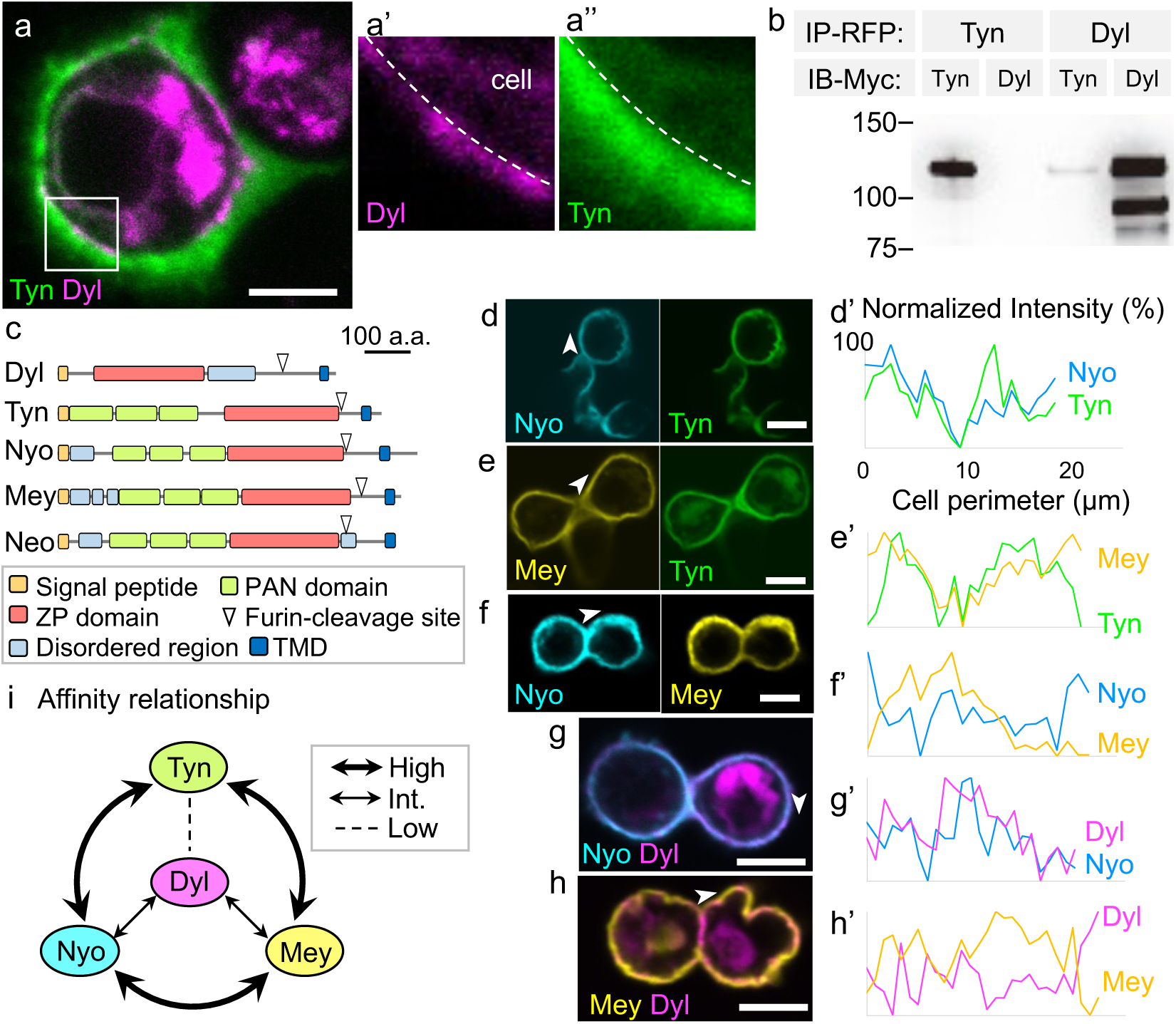
Self organization of ZPD matrices in cultured cells. **a,** A two-layered ECM structure on S2 cells expressing Tyn (green) and Dyl (magenta). **a’, a’’** Enlarged views of the boxed region in **a.** Dotted line indicates cell membrane. **b,** Immunoprecipitation assays. Tyn and Dyl constructs tagged with RFP or Myc were separately transfected to S2 cells. Cells were mixed in pairwise manners. Cell extracts immunoprecipitated using anti-RFP antibody were probed with anti-Myc antibody. Homo-oligomers of Tyn and Dyl were abundantly detected. A weak band of RFP-Dyl and Myc-Tyn complex was also detected. No signal was detected from the reciprocal combination of RFP-Tyn and Myc-Dyl, due to higher expression level of the Tyn construct. **c,** Structures of the five ZPD proteins. **d-h,** Pairwise expression of Tyn, Dyl, Nyo and Mey. **d’-h’,** Fluorescence intensity profiles of ZPD protein signals in the cell perimeter starting from the points shown by arrowhead. **i,** A diagram of the affinity relationship of ZPD proteins inferred from their mixing patterns. Scale bar: 5 μm.

Live imaging of S2 cells expressing the Tyn matrix showed the rapidly changing shape of the matrix and deformation of the shape of actively moving cells (Tyn, Supplementary video 1). Dyl matrix was also deformed and was split upon cell division (Dyl, Supplementary video 2). Tyn matrix covering the cell surface expanded and was ruptured during cell mass increase (Supplementary video 3 and Extended Data Fig. 2a). A part of the cell cortex pinched by Tyn matrix accumulated F-actin, suggesting high mechanical stress (Extended Data Fig. 2b). These data indicate Tyn matrix, and possibly Dyl matrix, are elastic materials with a rigidity sufficient for deforming the interphase cell cortex (Young’s modulus ∼1.0 kPa^26^).

Outside of the ZPD, Dyl has a disorganized region of flexible conformations, and Tyn has three plasminogen N-terminal (PAN) domains (Fig. 2c). Mutant proteins of Dyl without the disordered region (Dyl^ZP^) and Tyn without any PAN domains (Tyn^ZP^) formed ECM and those lacking ZP domains (Dyl^ΔZP^, Tyn^ΔZP^) failed to be integrated into ECM (Extended Data Fig. 2c-f). Cleavage of the signal peptide and the furin-dependent C-terminal transmembrane region of Dyl^ZP^ and Tyn^ZP^ leaves ZPD only, suggesting ZPD is necessary and sufficient for ECM formation. Co-expression of Tyn and Dyl^ZP^, and Tyn^ZP^ and Dly formed distinct sorting patterns similar to those of the full-length Tyn-Dyl pair (Extended Data Fig. 2g, h and Fig. 2a). These data indicate that Tyn and Dyl self-oligomerize and sort into distinct ECMs through the function of their ZPDs. Importantly, the sorted patterns of Tyn and Dyl resembled the patterns of layer II–III cloud ECM and layer I, respectively, in the olf.

Pairwise expression of Tyn, Nyo, and Mey, or expression of all three formed completely mixed ECMs (Fig. 2d-f, d’-f’, Extended Data Fig. 2i). On the other hand, co-expressed Nyo and Dyl, and Mey and Dyl formed partially separated ECMs (Fig. 2g, h). Tyn^ZP^ also mixed well with Nyo and Mey, and Dyl^ZP^ matrix sorted partially with Nyo and Mey (Extended Data Fig. 2j-m). The results indicate that ECM ZPD proteins have distinct mutual affinities: the Tyn matrix has a high affinity for Nyo and Mey matrices, while its affinity for the Dyl matrix is low. Nyo and Mey matrices have an intermediate affinity for the Dyl matrix. Collectively, these results indicate that ZPDs of Tyn and Dyl are responsible for sorting and mixing patterns with other ZPD proteins. Together with differential expression and sequential secretion in different cells (Extended Data Fig. 1f-j), cell-specific compartmentalized ECM patterns in the olf, mech, and sp are established.

### Dyl organizes envelope formation

We next addressed the role of Dyl in cuticle formation. Knockdown of Dyl in the sensory organ (with *neur-gal4*) caused severe loss of those organs in the adult maxillary palp (Fig. 3a, b and Extended Data Fig. 3a, b). In the olf at 44 h APF, the total amount of envelope was reduced (Fig. 3c, d, envelope coverage, control: 96.8% and 97.7% n=2, dyl RNAi line #1, 67.5% and 32.8%, n=2, line #2 68.2%, n=1). On the mech, the envelope was selectively lost in the region without the actin bundle (Fig. 3e, f), where normally Dyl is enriched as longitudinal stripes, as previously shown for the mech in the notum^20^ (Fig. 3g, h).

**Figure 3.**
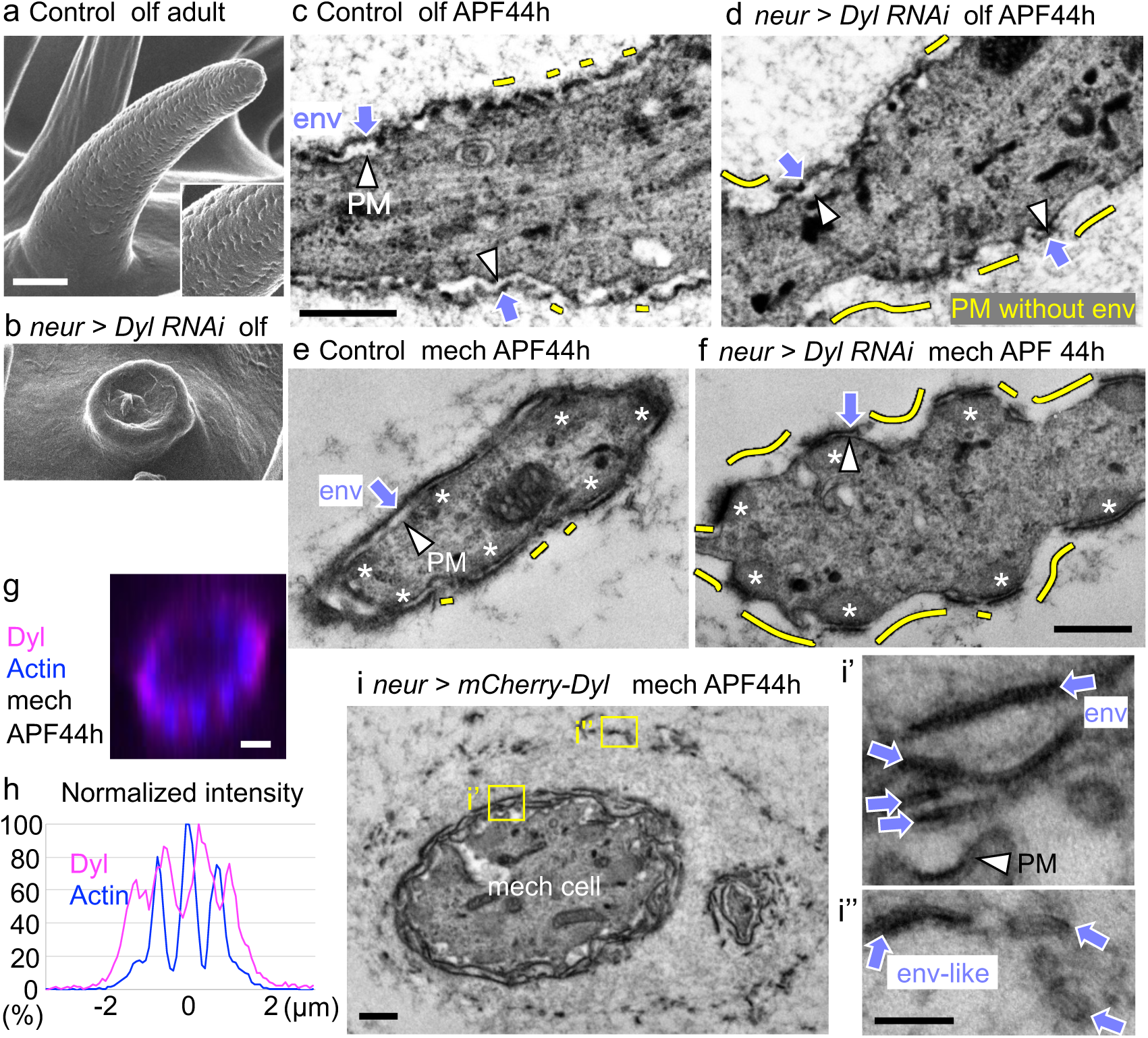
Dyl is required for envelope formation. **a, b,** Olfactory bristles of normal (**a**) and Dyl-knocked-down (**b**) adult flies. **c, d,** Longitudinal TEM views of olf at 44 h APF. In the control (**c**), the plasma membrane (PM, arrowheads) is covered by the envelope (env, arrows) except for small gaps (yellow lines). **d,** Dyl RNAi caused a dramatic loss of env (yellow lines). **e, f,** Cross-sectional views of mechanosensory bristles in control (**e**) and Dyl-knocked-down flies (**f**). A dramatic loss of the envelope by Dyl suppression (yellow lines) occurred in the region free of actin bundles (asterisks). **g,** A fluorescent image of Dyl and actin in a cross-section of mech. **h,** The signal intensity profiles of Dyl and actin. **i,** The multilayered env formation induced by Dyl overexpression in mech. **i’, i’’,** Enlargements of the boxed regions. Scale bars: 1 μm for **a, b,** and **g,** 100 nm for **i’** and **i’’,** 500 nm for the others.

Overexpression of Dyl caused malformation and nanopore loss in the olf (Fig. 5f-h, details will be described later), and truncation of the mech (Extended Data Fig. 3c). In the mech at 44 h APF, overexpressed Dyl disrupted the endogenous striped pattern of Dyl, instead forming thick, stratified ECM layers that covered the distal half of the cell (Extended Data Fig. 3d, d’, d’’). At the corresponding site, multiple envelope-like layers surrounded the cell surface, extending up to 1 μm from the cell surface (Fig. 3i, i’, i’’). The result indicates that Dyl plays an essential role in envelope formation around a subset of the plasma membrane. Its activity to induce multiple envelope-like structures indicates Dyl has a key organizing role in envelope formation.

### Cloud ECM ensures regular nanopore formation

To understand the role of cloud ECM, we individually knocked-down its components Tyn, Mey, Nyo, and Neo. Tyn was lost around olf bristles, indicating that Tyn in cloud ECM is synthesized cell-autonomously (Extended Data Fig. 4a). However, cloud ECM was still observed under transmission EM, and Mey, Nyo, and Neo were present around the olf hairs (Extended Data Fig. 4b, c). Those olf hair cells at 44 h APF increased width by 25% and length by 27% (Extended Data Fig. 4d Tyn RNAi-1), suggesting Tyn has a function to restrict the rapid growth of the olf around 44h APF (Extended Fig. 4e). However, the resultant adult olf cuticle still formed nanopores. Similarly, the appearance of Mey, Nyo, or Neo knock-down on the adult olf cuticle was indistinguishable from control. The results indicate that cloud ECM is composed of multiple protein components and the knockdown of individual ZPD proteins was insufficient to eliminate cloud ECM in toto.

**Figure 4.**
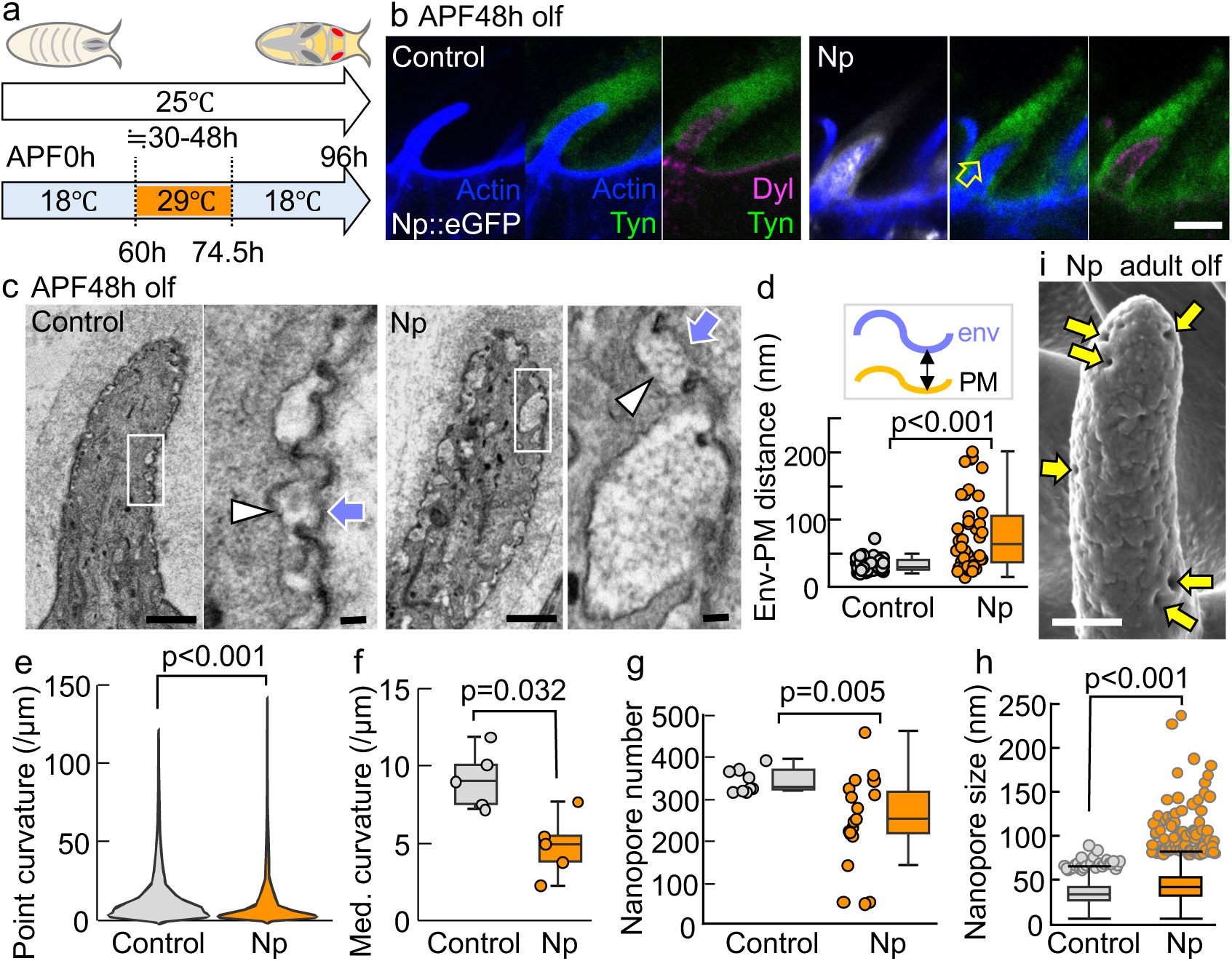
ZPD protein degradation impaired nanopore formation. **a,** The temperature shift protocol of the Np overexpression experiment. **b,** Olfactory bristles at 48 h APF in the control (left panels) and Np overexpression groups (right panels). The arrow indicates the gap between Tyn and the cell. Scale bar: 5 μm. **c,** Longitudinal sections of olfactory hairs by EM. The right panel (scale bar: 100 nm) is an enlarged view of the highlighted area in the left panel (scale bar: 1 μm) for each. Arrowhead: PM. Arrow: env. **d,** The distances between env and PM was measured at concave parts (Control: n=42 and Np: n=41 concave parts from 5 hairs in a fly). **e, f,** Point curvature measured using the Kappa plugin in Fiji (Control: n=39552 points, Np: n=27520 points) and the median curvature for each hair (n=5 hairs from a fly for each group). **g, h,** Quantification of nanopore number (Control: n=9 hairs from 3 flies, Np: n=20 hairs from 7 flies) and size (Control: n=1550 pores from 3 flies, Np: n=2493 pores from 7 flies, boxplots with data points for the outliers were displayed) based on SEM images. **i,** SEM view of an olfactory bristle with Np overexpression. Arrows indicate enlarged pores. Scale bar: 1 μm. Mann-Whitney U test was used for all statistical analyses.

As an alternative approach, we attempted to remove multiple ECM components by targeting protease expression. Among Stubble (Sb), Lumens interrupted (Lint), Notopleural (Np), and Tracheal-prostasin (Tpr), that are known to degrade ZPD proteins Dumpy (Dpy) and Piopio (Pio)^27,28^, Np, a homolog of Human matriptase, caused recognizable defects in the olf. Np::eGFP was expressed by *neur-gal4* during the time window corresponding to 30 to 48 h APF at 25°C to avoid the lethality by its constitutive expression (Fig. 4a).

Np caused cleavage of Mey and Nyo, but not Tyn, in S2 cells (Extended Data Fig. 4f). Although the Tyn level was unchanged in the olf hairs as well, the distance of Tyn from the cell surface was expanded in Np expressing olf (Fig. 4b, Extended Data Fig. 4g). Dyl levels remained unchanged (Fig. 4b). We noted with ∼55% increase in the olf width (Fig. 4b, Extended Data Fig. 4h). EM views of Np-expressing olf cells showed that the distance between the envelope and plasma membrane was significantly increased (Fig. 4c, d). The envelope took a partially wavy shape, but the curvature of the envelope decreased (Fig. 4e, f). The Np expression did not compromise the growth of the olf and mech (Extended Data Fig. 4i). Remarkably however, the number of nanopores decreased, and the size became irregular, with large pores being observed (Fig. 4g-i). Those results indicate that Np expression removes a subset of layer II ECM components, permitting overexpansion of the olf cell diameter and detachment of envelope from the cell membrane. Thus, cloud ECM provides the compressive force required for regular nanopore formation.

### Envelope-membrane proximity is essential for nanopore formation

In the olf at 42-44h APF, overexpressed Dyl formed tight coverage of the shaft cells and caused detachment of the envelope from the cell membrane (Fig. 5b, c), as observed in Np overexpression (Fig. 4c,d). Moreover, the envelope of Dyl-overexpressing olf often exhibited a flat appearance (Fig. 5b, d, e). Consistently, the number of nanopores decreased and larger pores were observed (Fig. 5f-h). The result indicates that the excess Dyl caused the displacement of the envelope from the plasma membrane and the loss of nanopore.

**Figure 5.**
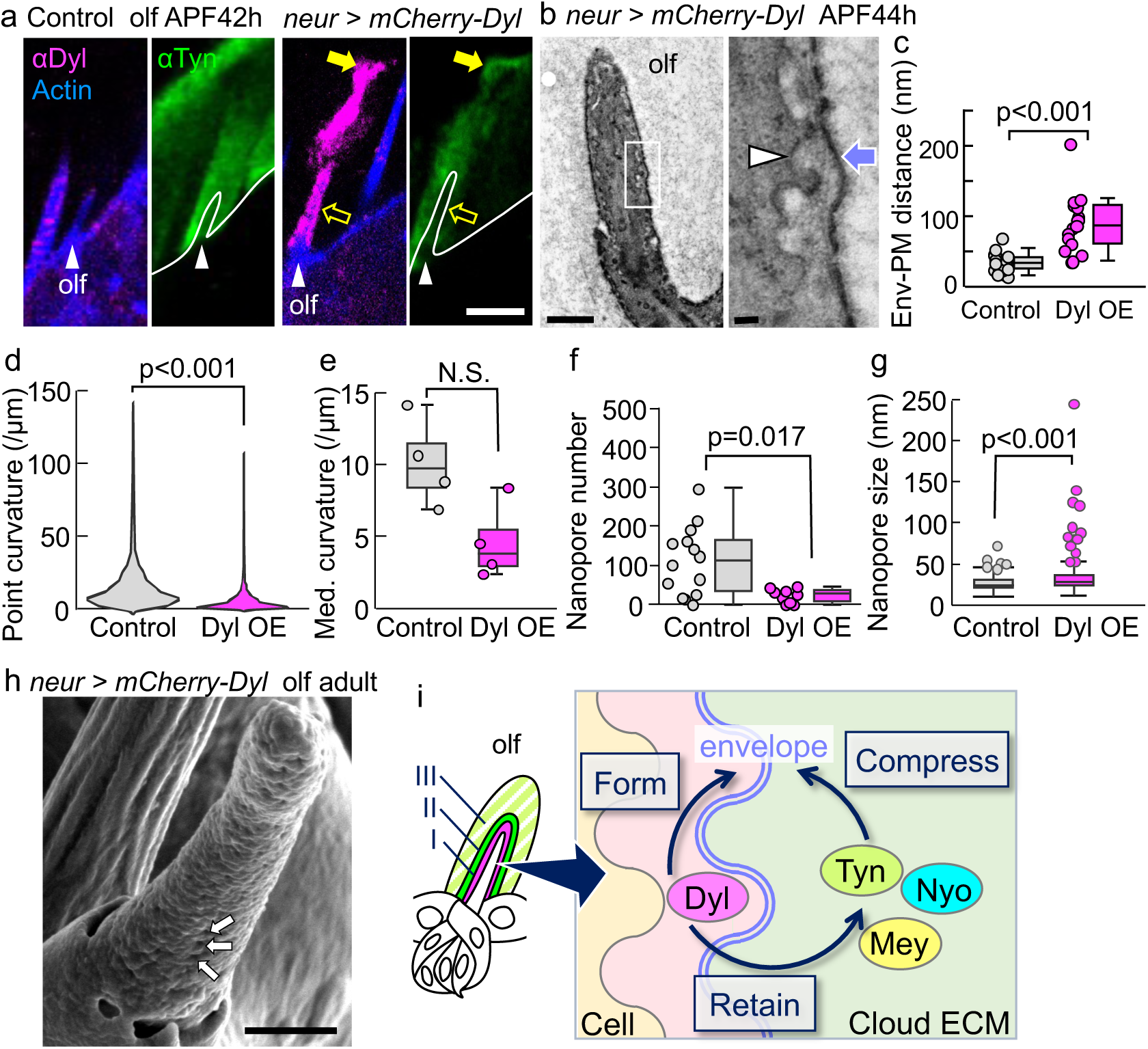
The envelope-plasma membrane proximity controls nanopore formation. **a,** Overexpression of Dyl in the olf hair cells. Dyl, Tyn and F-actin staining of control (left) and Dyl-overexpressing (Dyl-OE, right) olf hair cells (arrowheads). Overexpressed Dyl covered the olf cell surface (open arrow) and accumulated at the tip (arrow), which carried endogenous Tyn away. Scale bar: 5 μm. **b,** An electron micrograph of Dyl-OE olfactory hair cell (left, Scale bar: 1 μm). The enlargement of the boxed area (right) shows a wavy plasma membrane (arrowhead) and a relatively straight envelope (arrow). Scale bar: 100 nm. Note that electron-dense concave parts of the envelope were absent in this image. **c,** The distances between the envelope and plasma membrane were measured in the TEM images (Control: n=19 and Dyl-OE: n=16 from 2 hairs in 2 flies for each group). **d, e,** Point curvature (Control: n=14080 points, Dyl-OE: n=11520 points) and the median curvature for each hair (n=4 hairs from two flies for each group, p=0.057). **f, g,** Adult cuticle of Dyl-OE olfactory hair showed reduced nanopores (Control: n=14 hairs from 5 flies, Dyl-OE: n=9 hairs from 3 flies), and irregular nanopore size (Control: n=790 pores from 3 flies, Dyl-OE: n=108 pores from 7 flies, boxplots with data points for the outliers were displayed). **h,** SEM view of a Dyl-OE olfactory bristle. Arrow: normal nanopore. Scale bar: 1 μm. **i,** A model of envelope formation and patterning by ZPD proteins. I, II, and III surrounding the olfactory hair indicate the layers I: Dyl, II: Tyn, Nyo, and Mey, III: Tyn, Mey, and Neo, respectively (also shown in Extended Data Fig. 1j). In this working hypothesis, Dyl retains cloud ECM outside and promotes envelope formation at the interface between layer I and II. layers II and III (cloud ECM) compress the envelope so that it tracks the wavy shape of the plasma membrane. Mann-Whitney U test was used for all statistical analyses.

## Discussion

In this work, we tested the hypothesis that the curved envelope of olf is formed by tracking the undulated plasma membrane^3^. In support of this model, disruption of the close association between the envelope and plasma membrane by reducing the compressive force of layer II/III cloud ECM or thickening of layer I ECM caused loss or malformation of the nanopore. Apical ECM of insects undergoes rapid deposition and replacement during cuticle development^29–31^. The ZPD proteins Dyl, Tyn, Nyo, and Mey form rapidly changing and cell-type-specific ECM distributions during the early phase of sensory organ development, prior to the cuticle deposition. The functions of the ZPD aECM are twofold. Layer I occupies a thin territory of up to 125 nm above the plasma membrane; the envelope forms at the outer border of layer I. The self-organizing activity of Dyl to form a layered ECM structure in cultured cells may be responsible for layer I formation. The surface of the Dyl ECM layer sorts Tyn to the outside to form a two-layered organization. The prospective envelope components may be segregated from the Dyl layer and are assembled at the interface to construct the envelope in such a way that the amphipathic molecules align at the water-oil interface.

The second role of ZPD ECM is the formation of cloud ECM that occupies layer II and III of the olf. Proteolysis by Np caused the widening of the olf, suggesting that the ZPD ECM containing Tyn, Nyo, Mey, and possibly other proteins applies compressive force essential for nanopore formation. The high mixing activity of Tyn, Mey, and Nyo indicates that cloud ECM is a composite of multiple ZPD and other molecules. Removal of any one of those ZPD proteins did not cause a significant change in the cuticle secretion and nanopore formation, indicating the multiple molecular compositions of cloud ECM makes the matrix robust against the fluctuation of ZPD protein quantity. The expression of transient ZPD matrices prior to the cuticle formation has been described as “precuticle” during vulval and sensory organ development in *C. elegans*^32,33^. Thus, the mechanical role of ZPD matrices in cuticle patterning reported here may play similar roles in a wide range of animal species.

The plasma membrane of the olf cells is highly undulated, whereas the plasma membrane in the mech and sp is flat (Extended Data Fig. 4j, j’’, k). The difference in the membrane shape corresponds to the presence of actin bundles underlying the membrane. In the mech, membranes attached to the actin bundles are flat, while those in the inter-bundle region are abundant with membrane protrusions (plasma membrane plaque, Extended Data Fig. 4j’) and often convoluted^34^. Actin bundles in the trachea have been shown to suppress endocytosis^35^. Low abundance of subcortical actin in the olf cells may thus permit frequent membrane deformation under abundant membrane supply, due to elevated ER activity in olf (Inagaki et al., submitted), while the undulated membrane serves as a template to shape the envelope under compressive force into a curved structure. In this condition of rapid remodeling of the plasma membrane, Dyl is essential for stabilizing the envelope formation.

ZPD proteins form various configurations matrix. Dpy forms tensile filaments that anchor a part of imaginal disc to direct tissue flow^36–40^ and restrict tube elongation by forming luminal filaments in the trachea^41^. Tyn, Nyo, and Mey form a 3-dimensional cloud ECM covering the cells to set up a compressive mechanical environment, and Dyl forms a sheet that organizes layer formation of the envelope. In cultured cells, ZPD of Dyl, Tyn, and Dpy are, in principle, sufficient for directing molecular assembly into the sheet and cloud (this study), and filamentous forms^39^. Further investigations into ZPD structures may lead to an improved understanding of principles underlying modular matrix design based on the ZPD. The findings presented here study will pave the way for uncovering various morphogenetic mechanisms and the development of novel biomimetic technologies.

## Methods

### Fly strains

*Drosophila melanogaster* strains were cultured with standard yeast-cornmeal-agar food at 25℃ otherwise noted. For staging the pupae, we collected white pupae at APF 0 h, allowed them to grow for a specific time, and then fixed them. *w[*]; P{UAS-Np::eGFP}* is a gift from M. Behr^27,28^.

### Temperature control for Np overexpression experiment

Parental flies were maintained at 18℃, and offspring at the white pupal stage were collected. After 60 h, the pupae were transferred to 29℃ for 14.5 h and then returned to 18℃. The heating time corresponds to around 30-48 h APF period within the total 96-h pupal stage at 25℃.

### Generation of expression vectors and transgenic flies

For the construction of expression vectors, the coding sequence of *dyl* or *tyn* with a tag right after the signal peptide was introduced into pUASTattB plasmids. These plasmids were used as PCR templates for site-directed mutagenesis with the following primers, each paired with its reverse complement primer: “CAAGCACCTGCAGGTGGAGTACGGTCTGCCC” for pUASTattB-mCherry-Dyl^ΔZP^, “CCCCAGATTGGAGCCAAGGATGACCTGAGTGC” for pUASTattB-mCherry-Dyl^ZP^, “GCTATGATGTTTCTGTTGAATCTTTGGGGAGACGG” for pUASTattB-mCherry-Tyn^ΔZP^, and “GGAAGCGGAGGTTCCCCACTGATCGATCATGG” for pUASTattB-mCherry-Tyn^ZP^. pUASTattB-mVenus-Nyo and pUASTattB-HA-Mey were constructed and packaged by VectorBuilder (vectorbuilder.com). pUASTattB-mCherry-Dyl or pUASTattB-mCherry-Tyn (150ng/μl each) were injected into the *y, v, nos-φC31; attP40* strain and the UAS strains were established.

### Generation of knock-in flies by genome editing

mVenus or mScarlet sandwiched with linkers were inserted after the sequence of signal peptide (SP) in *dyl* or *tyn* gene with the use of the Scarless gene editing system (https://flycrispr.org/scarless-gene-editing), a combination of CRISPR/Cas9-mediated homology-directed repair and precise excision by PBac transposase as follows. The gRNAs for Dyl (GCCCATTTACGGTGCGCCCC) and Tyn (GTGATCGATCTTCGATGCAC) close to the insertion sites were designed with CRISPR Target Finder^42^ (http://targetfinder.flycrispr.neuro.brown.edu) and cloned into pBFv-U6.2 vector^43^ as described in our precious work^24^. To generate donor plasmids, sfGFP in pHD-sfGFP-ScarlessDsRed (Addgene #80811) was first replaced by a fluorescent tag mVenus or mScarlet. A continuous genomic region, including the sequence from downstream of the SP to the gRNA site and its homology arms more than 500 bp for both sides, was amplified and inserted into a backbone vector. Finally, the marker cassette flanked by transposon ends was inserted right after the SP in the modified backbone vector. Importantly, silent mutations were introduced into the gRNA sequence of the donor plasmid to avoid mutagenesis after the knock-in. The donor and gRNA constructs (150ng/μl each) were injected into the *y[2] cho[2] v[1]; attP40{nos-Cas9}/CyO* strain. Once the tag was introduced into the genome, the DsRed marker was removed by crossing with *w[1118]; CyO, P{Tub-PBac\T}2/wg[Sp-1]; l(3)*[*]/TM6B, Tb[1]* and fly strains were established.

### Fluorescence microscopy

Pupae were removed from the pupal case and gently scratched to facilitate fixation, then immediately transferred to the fixation buffer (4% formaldehyde in phosphate-buffered saline, PBS). After a few hours or one night at 4℃, pupal cuticles were removed, and collected head tissues were washed three times with 2% bovine serum albumin (BSA), 0.2% of Triton X-100 in PBS, followed by antibody incubation. The head tissue underwent further dissection to isolate the maxillary palps with labellum and mounted with VECTASHIELD Mounting Medium (VECTOR LABORATORIES H-1000 or H-2000). Images were taken by using Olympus FV1000 with UPLSAPO60XW 60x NA-1.2 water-immersion lens. Rat anti-Dyl and rabbit anti-Tyn antibodies (1:300) were a gift from F. Payre^19^. Peptides Cys-VATPNGSELPKPLS-OH for anti-Mey, Cys-VKQTNPRTNVTPSP-OH for anti-Nyo, and Cys-SLGSEEDSIYYDNA-OH for anti-Neo were used to generate affinity-purified antibodies (1:500) using custom antibody service provided by SCRUM Inc., Japan. Alexa Fluor^TM^ Plus 405 Phalloidin (Thermo Fisher Scientific, Cat# A30104) was used for actin staining.

### Transmission electron microscopy

The sample preparation was done following a previous paper^3^ with some modifications. Pupal head tissues were collected as described above, in a different fixative solution with 2.5% glutaraldehyde, 2% formaldehyde in 0.1 M cacodylate buffer, pH7.4 for over one night but within one week. The tissues were washed for 10 min in 0.1 M cacodylate buffer three times and post-fixed in 2% OsO_4_, 1.5% (wt/vol) potassium ferrocyanide, 2 mM CaCl_2_, and 0.15 M sodium cacodylate buffer (pH 7.4) for 2 h on ice with light shielding. The tissues were washed for 10 min in water three times, stained with 0.5% uranyl acetate at 4℃ overnight, and washed in water for 15 min twice, followed by dehydration in a series of graded ethanol. The ethanol was replaced by propylene oxide and then resin. After resin polymerization at 60℃ for more than 48 hours, 60 nm-thickness ultrathin sections were obtained and observed by JEM-1400Plus (JEOL). The following two experiments were conducted by S. I. with modifications, and the data was from antennal large basiconic sensilla, which we consider are analogous to the olfactory bristles found in maxillary palp. First, for stronger staining of the envelope (Fig. 1d), the post-fixation was followed by washes, 1% thiocarbohydrazide (TCH) treatment for 1 h at 60℃, additional washes, and the second post-fixation 1 h on ice. Second, for ruthenium red (RR) staining, 0.5 mg/ml RR (SIGMA 00541, dissolved in water) was added to the post-fixation solution.

### Immuno-electron microscopy

Pupae were immersed in a 4% PFA solution in PBS at 4°C for a few hours or overnight, and the isolated head tissues were then washed twice with PBS for 10 minutes each and treated with a solution of 1% BSA and 0.01% Triton X-100 in PBS for 10 minutes. Overnight incubation with a primary anti-GFP rabbit antibody (1:200, MBL Cat# 598) in the permeabilization solution followed. The tissues underwent three 10-minute washes and the application of a secondary goat anti-rabbit Fab’ Alexa Fluor 594 FluoroNanogold (1:200, NAN Cat# 7304) for 2 hours at room temperature. After the wash with the permeabilization solution for 10 minutes and two 10-minute washes in PBS, post-fixation was performed using 0.5% glutaraldehyde in PBS for 30 minutes at 4°C. After washing the tissues twice in water for 10 minutes each, a gold enhancement with GoldEnhance^TM^ EM Plus (NAN, Cat# 2114) was conducted for 5 minutes. Following another rinse in water, the tissues were dehydrated and embedded in resin, the same as the transmission EM protocol described above. The ultrathin sections on grids underwent double staining using uranyl acetate and lead citrate, followed by washing and observation with JEM-1400Plus (JEOL).

### Electron microscopy for scanning cuticle surface structure

Field emission scanning electron microscopy (FE-SEM) and helium ion microscopy (HIM) were performed based on a previous report^3^. Adult flies were collected in 70% ethanol and stored at room temperature. Head tissues were fixed and dehydrated as described for the transmission EM analysis but without uranyl acetate treatment. In the middle of this study, cacodylate buffer was replaced with PBS as it yielded the same results. For drying, the samples were placed in a desiccator for about a week. Finally, the samples were coated with osmium at 5 nm twice (Tennant 20, Meiwafosis) and observed by FE-SEM (JSM-IT700HR, JEOL) or HIM (ORION Plus, Carl Zeiss at the Nano-processing facility in AIST Tsukuba, Japan).

### Cell culture, transfection, staining, and observation

*Drosophila* S2 cells (RIKEN Bioresource Center, RCB1153; RRID: CVCL_Z232^44^) were cultured at 25°C in Schneider’s *Drosophila* medium (Thermo Fisher/Gibco, Cat# 21720024) with 10% fetal bovine serum or in Sf-900 II SFM (Thermo Fisher/Gibco, Cat# 10902096), both supplemented with Penicillin Streptomycin (Gibco Pen Strep #15140-122) at a dilution of 1:100. Transfection was performed using TransIT-Insect Transfection Reagent (Mirus Bio, Cat# MIR6104), following the protocol provided but with a reduced amount of each DNA construct (50-200 ng, pWAGal4 and pUASTattB plasmids described above). Cultures were observed after 2-3 days. The staining of transfected cells was performed on coverslips submerged in the medium during culture. Alternatively, transfected cells were dropped onto coverslips and allowed to adhere for 30-60 minutes before the staining. The cells were washed with PBS and fixed with 4% paraformaldehyde for 15 minutes at room temperature. The cells were washed again with PBS. For intracellular staining, cells were treated with a blocking solution (BW: 0.2% Triton X-100, 0.2% Tween-20, 0.1% BSA in PBS) for 15 minutes. Primary and secondary antibodies were applied sequentially, each followed by three PBS washes. Antibodies used were rabbit anti-Myc pAb (MBL, Code 562), rat anti-HA (Roche, 3F10), Alexa Fluor^TM^ Plus 405 Phalloidin (Thermo Fisher Scientific, Cat# A30104), goat anti-rabbit IgG Alexa Fluor Plus 405 (Invitrogen), and goat anti-rat-IgG-StarRed (Abberior STRED-1007), all used at a dilution of 1:200. After staining, the cells were mounted and observed using an Olympus FV1000 microscope.

### Co-immunoprecipitation and Western Blot Analysis

S2 cells harboring a transgene pMT-mCherry-Tyn, pMT-mCherry-Dyl, pMT-MBP-Myc-Tyn, or pMT-MBP-Myc-Dyl were generated by co-transfection of pCopHygro and each pMT plasmid constructed using pMT/V5-HisA (Invirogen V412020) followed by selection in 300 μg/ml Hygromycin B-containing medium (FUJIFILM, Cat# 084-07681). A pair of transgenic cell lines at a density of 5 × 10^5^ cells/ml each were combined in 25 cm^2^ flasks, in a total of 5 ml of media. Expression of the constructs was induced with 500 μM CuSO_4_ solution. Four days post-induction, when cells reached confluence, supernatant was collected and centrifuged at 1000 rpm. The supernatant was further centrifuged at 19,500 g. The supernatant was mixed with 50 μL of anti-RFP or anti-Myc magnetic beads and incubated at 4°C for 1 h with rotation. Beads were washed four times with 50 mM Tris-HCl (pH 7.5) containing 150 mM NaCl and 0.05% NP-40. The proteins were eluted with 30 μl Laemmli SDS sample buffer and heated at 95°C for 5 minutes before loading onto SDS-PAGE gels. For Np cleavage assay, S2 cells were transfected with vectors UAS-mCherry-Tyn, UAS-mCherry-Dyl, UAS-HA-Mey, or UAS-Venus-Nyo alone or with UAS-Np::eGFP (a gift from M. Behr). Supernatants (15 μl each) and cell lysates (2 μl each prepared in RIPA buffer containing protease inhibitors) were analyzed by SDS-PAGE. Following SDS-PAGE, in both, proteins were transferred to PVDF membranes using an iBlot 2 Dry Blotting System. Membranes were blocked and incubated with primary antibodies diluted in iBind solution, following manufacturer’s recommendations. Detection was performed using Amersham ECL Prime Western Blotting Detection Reagent and visualized using the DavinchChemi chemiluminescence detection system. Antibodies used for the detection were anti-RFP-HRP-DirecT, anti-Myc-tag-HRP-DirecT, mouse monoclonal anti-HA-HRP-DirecT, rabbit anti-Nyo followed by anti-rabbit-IgG-HRP, and rabbit anti-GFP-HRP-DirecT (all from MBL, except for anti-Nyo generated in this study).

### Quantification and statistical analysis

ImageJ-Fiji (LOCI) processed and quantified images. JASP (JASP Team) performed statistical analysis and plot drawing.

## Acknowledgments

We are grateful to Matthias Behr, Isabelle Fernandes, and Francois Pyre for sharing flies and antibodies. We thank Vienna Drosophila Resource Center, Bloomington Drosophila Stock Center, and National Institute of Genetics for providing us with flies. We thank Takefumi Kondo and Alice Tsuboi for commenting on the manuscript. Shigenobu Yonemura and his laboratory members provided technical support for EM. HIM was conducted at the Nano-Processing Facility, supported by IBEC Innovation Platform, AIST.

## Author contributions

S.H. and Y.I. conceived the study. Y.I. performed the fly experiments, electron microscopy, and image data segmentation. H.W. performed cell culture experiments. S.I. performed a part of TEM analysis. S.H. and Y.I. analyzed the data and wrote the manuscript with input from all of the authors. All of the authors discussed the results and commented on the manuscript.

## Additional Information

Supplementary Information is available for this paper. Reprints and permissions information is available at www.nature.com/reprints. The image data that support the findings of this study are available in SSDB:repositry with the identifier doi: 10.24631/ssbd.repos.2024.07.370.

### Materials & Correspondence

Correspondence and requests for materials should be addressed to Shigeo Hayashi.

---

**Extended Data Figure 1.**
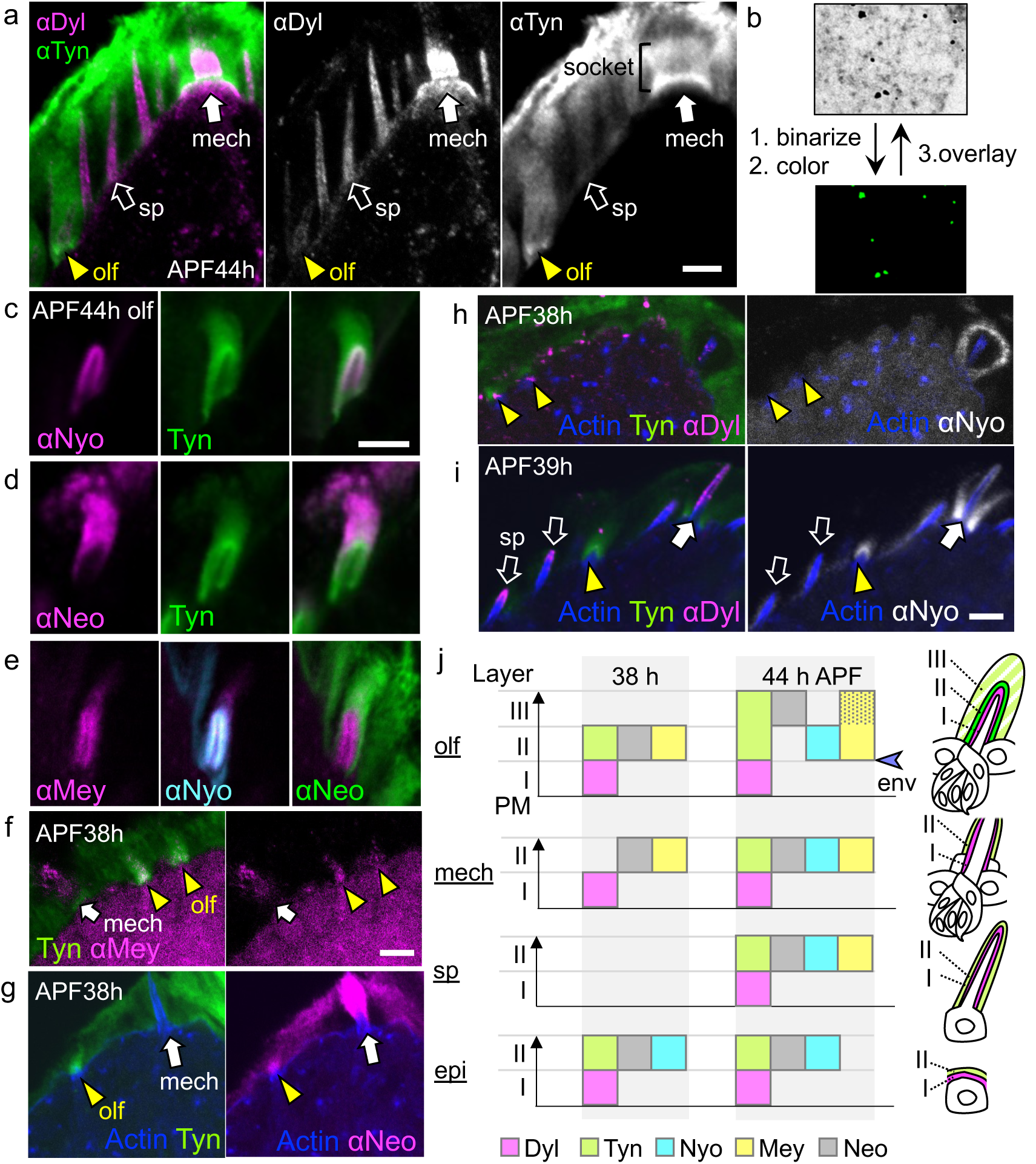
Cell-specific apical ECM structures. **a,** Distribution of Dyl and Tyn in the maxillary palp 44 h APF. Dyl is prominent on spinule (sp, open arrow) and mechanosensory hair cells (mech, arrow). Lower level of Dyl was also detected on the olfactory hair cell (olf, arrowhead) and epidermis. Tyn is abundant around the olf, the socket part of mech, and the exuvial space. **b,** Image processing of immuno-EM images. The electron-dense gold signals of Tyn were extracted by image binarization and coloration (green, lower panel) and then overlayed on the original image to create the image shown in Fig. 1i. At 44 h APF, Nyo (**c**) and Neo (**d**) are localized to the proximal and distal regions of the Tyn ECM, respectively. Mey is primarily located proximally, but a weak signal is also observed in the distal region (**e,** and a dotted area in **j**). **f-h,** At 38 h APF, Tyn, Mey, Neo, and Dyl are present on olf (yellow arrowheads), while Nyo is absent. Spinules have not grown yet in this stage. **i,** At 39 h APF, Nyo is observed at the proximal region (layer I) of olf. Spinules first appear at this stage with Dyl signals on their tip (open arrows). **j,** A diagram of the layered ECM structures on olf, mech, spinule hair cells and epidermis at 38 and 44 h APF. Scale bar: 5 μm.

**Extended Data Figure 2.**
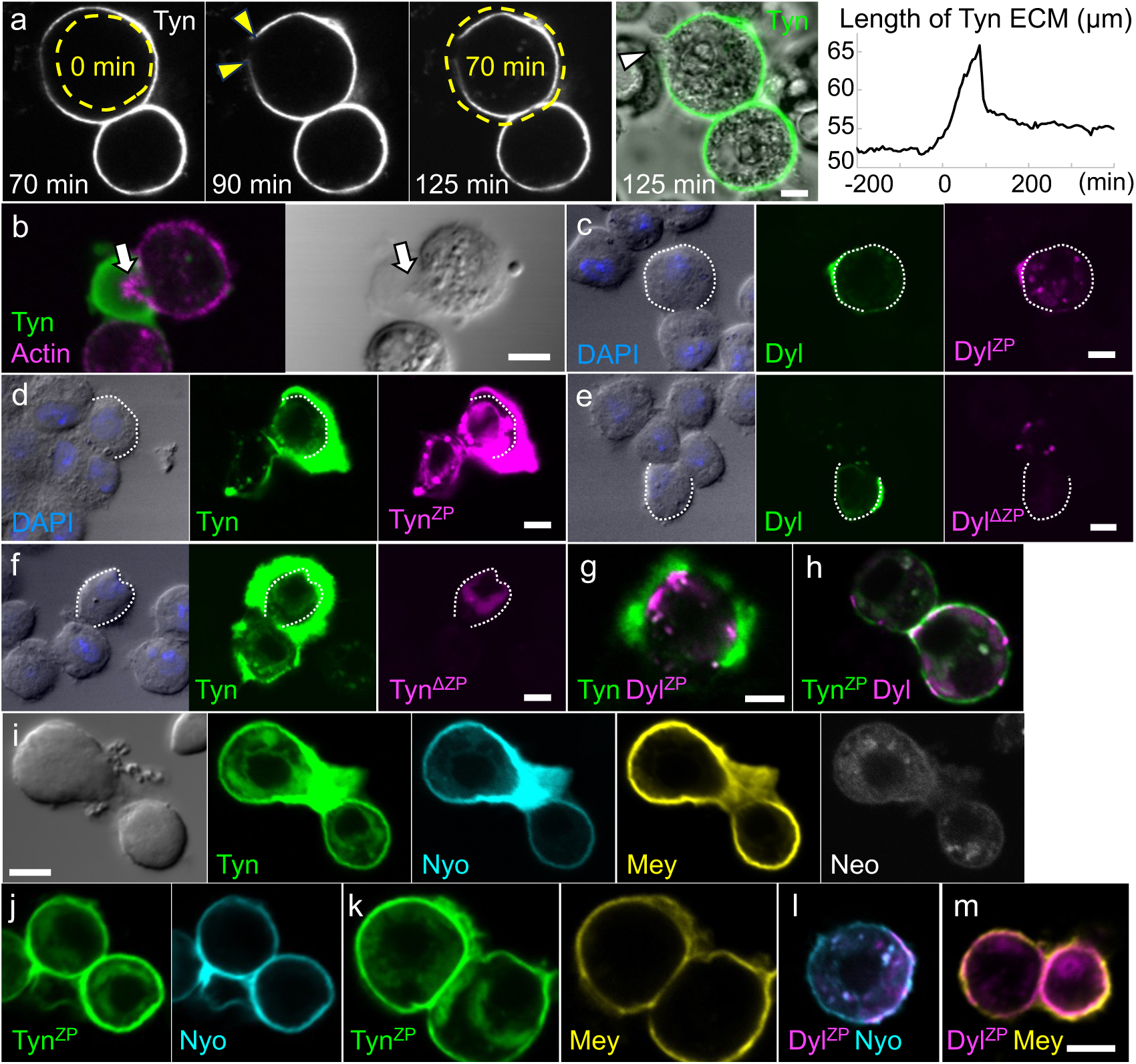
Elastic and mixing properties of ZPD matrices. **a,** Tyn matrix surrounding the upper S2 cell is stretched (70 min, as compared to the yellow dotted line showing the cell perimeter at 0 min, the starting point of Extended Data Video 3), punctured (90 min, yellow arrowheads), and shrunk (125 min, as compared to the cell perimeter at 70 min) during the cell volume increase. The cell membrane protruded from the broken ECM as indicated by the white arrowhead in DIC (Differential Interference Contrast) channel at 125 min. The length of Tyn ECM was shown in the rightmost panel. **b,** Tyn matrix deforms cell membrane (arrow). Note the high accumulation of F-actin in the protruded cell membrane in the Tyn matrix (arrow). **c-m,** Expression and mixing property of various forms of ZPD proteins. **c, d,** Myc-Dyl and mCherry-Dyl^ZP^ (Dyl without disordered region), and Myc-Tyn and mCherry-Tyn^ZP^ (Tyn without PAN domains) formed well-mixed ECM. **e, f,** Dyl^ΔZP^ and Tyn^ΔZP^ lacking ZPDs did not form ECM, even in the presence of their full-length forms. **g, h,** Combinations of Tyn and Dyl^ZP^, and Tyn^ZP^ and Dyl, formed segregated ECM as the combination of the full-length forms of each protein. **i,** Co-expressed mCherry-Tyn, mVenus-Nyo, and HA-Mey formed well-mixed ECM structure. Myc-Neo was not integrated into the ECM. **j-m,** Co-expressed with Nyo or Mey, Tyn^ZP^ and Dyl^ZP^ exhibited the same separation and polymerization behaviors as their full-length forms. Scale bars: 5 μm.

**Extended Data Figure 3.**
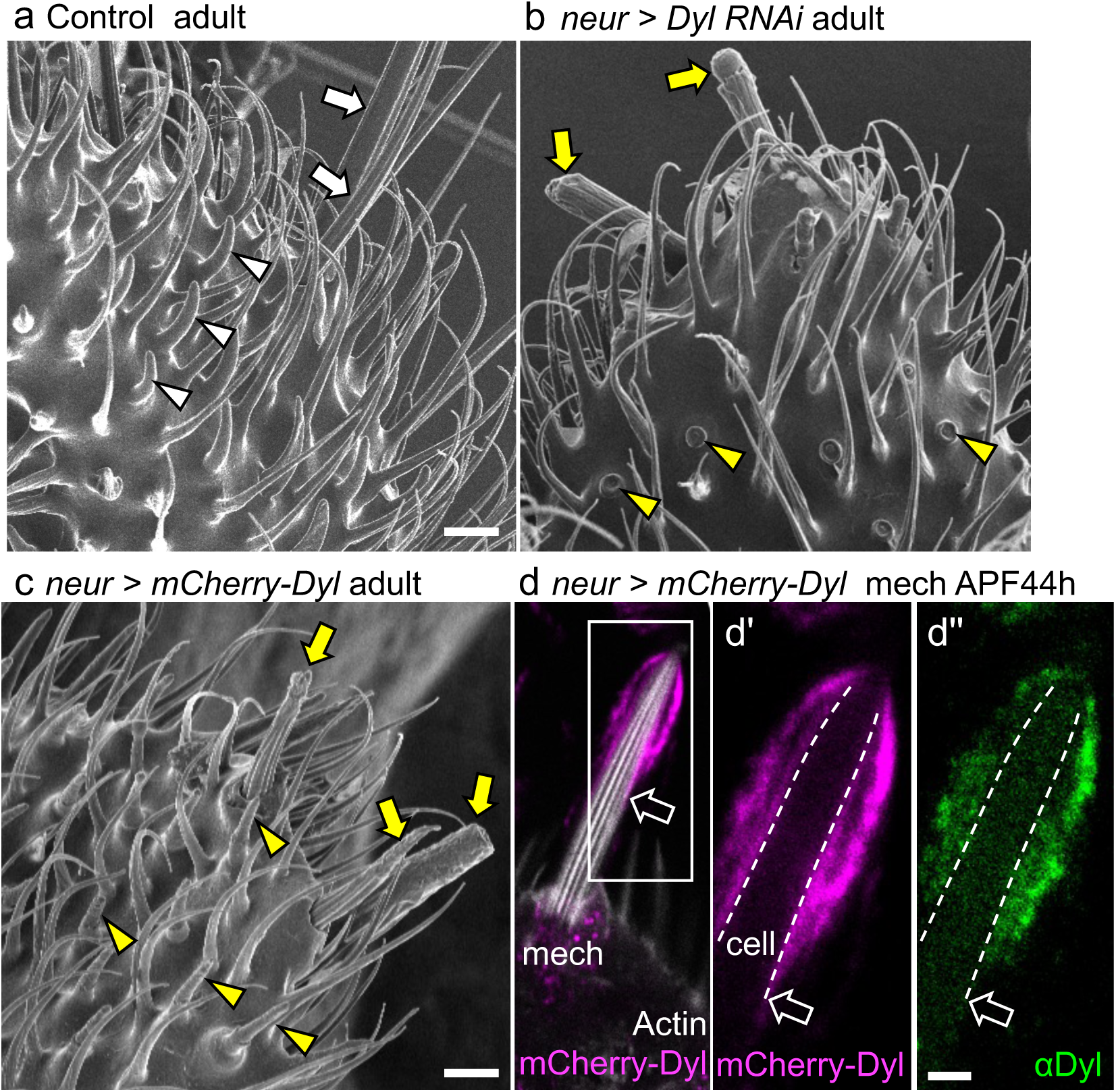
Roles of Dyl on bristle structures. **a-c,** Olfactory (arrowheads) and mechanosensory (arrows) bristles on the adult maxillary palp. **b,** Dyl RNAi driven by *neur-gal4* caused the shortening of mech (yellow arrows) and collapse of olf (yellow arrowheads, an enlarged view is shown in Fig. 3b). **c,** With mCherry-Dyl overexpression, many of the mechanosensory bristles showed shortening or disorganized surface structure (yellow arrows). The surface structure of the olf bristles (arrowheads) was disorganized, as shown in the enlarged view in Fig. 5h. **d,** Overexpressed mCherry-Dyl (magenta) accumulated around the distal part of the mechanosensory bristle (distal to the open arrow). The stripe pattern of actin bundles (white signal) is intact. mCherry-Dyl and immunolabeling of Dyl that labels both overexpressed and endogenous Dyl exhibited similar disorganized patterns (**d’, d’’**), indicating the endogenous stripe pattern of Dyl was disrupted. Scale bars: 5 μm.

**Extended Data Figure 4.**
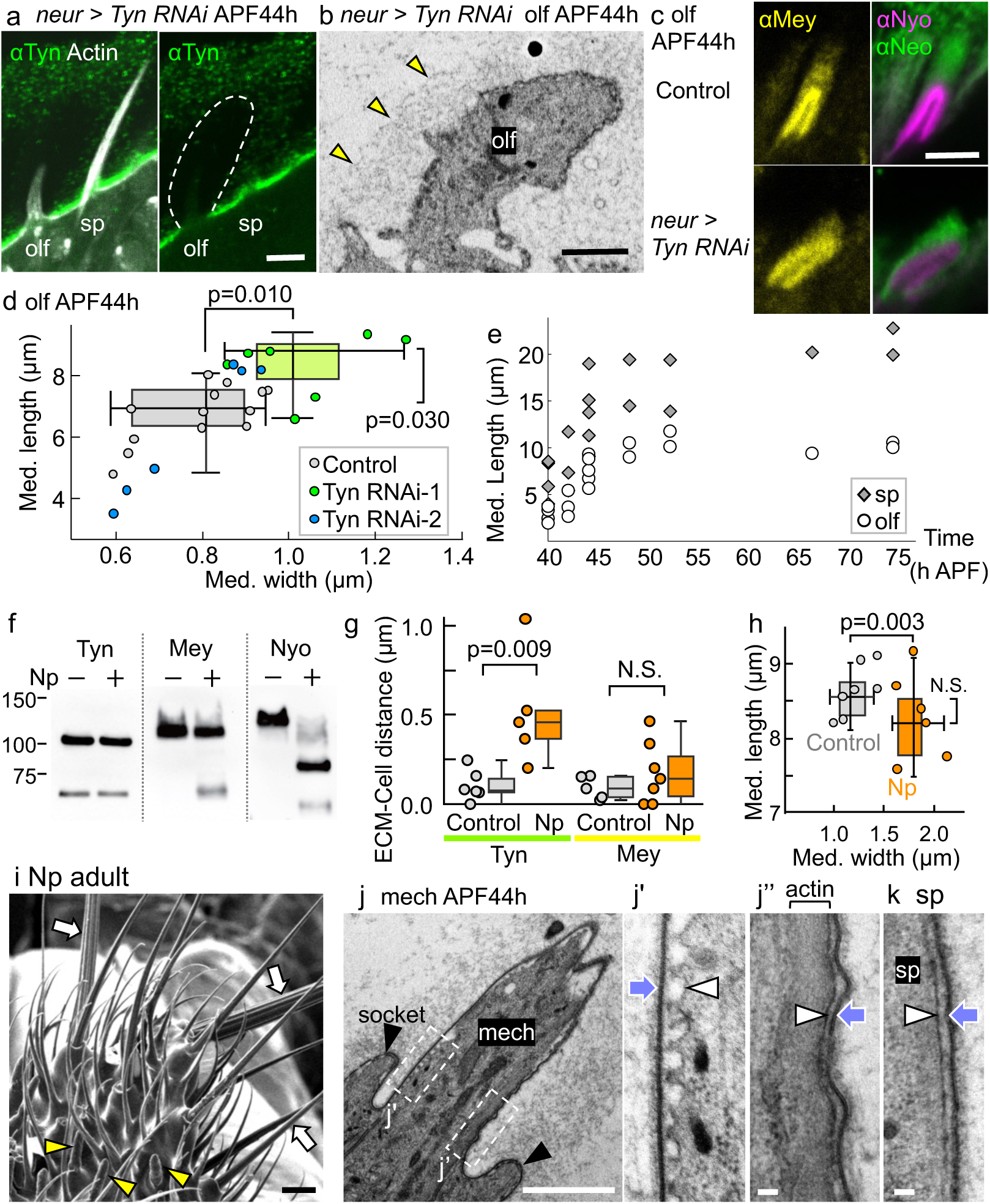
Supporting evidence for ECM-cell mechanical interactions in olfactory bristles. **a,** Tyn RNAi driven by *neur-gal4* caused a decrease of Tyn surrounding olf. The dotted line is presumed to be the boundary between the cloud ECM and the exuvial space. **b,** Despite the suppression of Tyn, cloud ECM was observed by TEM (arrowheads: outer boundary of cloud ECM). **c,** Mey, Nyo, and Neo are present in the layer II and III cloud ECM in the absence of Tyn. **d,** One of the two Tyn RNAi lines, Tyn-RNAi-1, induced the increased length and width (Kruskal-Wallis tests with Dunn’s post hoc and Bonferroni correction). Control: n=13, Tyn RNAi-1: n=7, Tyn RNAi-2: n=6 flies, the median value of 2 to 5 hairs from each fly. The box plot for Tyn RNAi-2 was not displayed because the variability was too high. **e,** The length of spinules and olfactory hairs from 40 to 75 h APF. Each data point represents a median value of 2 to 5 hairs from each fly. The cells elongated rapidly at 40-48 h. **f,** Western blot analysis of Tyn, Mey, and Nyo expressed in S2 cells with or without Np. **g,** Tyn-actin and Mey-actin distances were measured and presented as ECM-Cell distances. The difference in the Tyn-actin distances of the control and Np expressing groups was statistically tested (n=6 hairs from 2 flies for each group). Mey-actin distance also appears to have increased in the Np group, but statistical significance was not verified (Control: n=6 hairs from 2 flies, Np: n=8 hairs from 3 flies). Mann-Whitney U test. **h,** The width (horizontal axis) and length (vertical axis) of olf hair cells. The width but not the length increased by Np overexpression. Control: n=7, Np: n=6 flies, the median values of 3 to 5 hairs from each fly, Mann-Whitney U test. **i,** Np overexpression caused abnormality in olf (arrowheads, an enlarged hair shown in fig. 4i), while no obvious defect was observed in mech (arrows). **j,** TEM view of the control mech and the socket (black arrowheads). The plasma membrane (PM) of the region that is free of actin bundles showed a wavy shape (**j’**), while PM adjacent to the actin bundle was flat (**j’’**). In each case, the envelope was straight (arrows). **k,** Spinules, rich in actin bundles, showed straight PM and envelope. Arrowhead: PM, arrow: env. Scale bars: 5 μm for **a** and **c**, 1 μm for **b, i,** and **j,** 100 nm for **j’, j’’,** and **k.**

**Extended Data Video 1.** Constraint of cell shape and movement by expressed Tyn ECM. Note that a part of the cell covered by Tyn is constricted. S2 cells carrying the Myc-Tyn construct under the control of the metallothionein promoter were induced by the addition of 500 µM CuSO_4_. The recording started 160 min after induction. Tyn expression was detected with Alexa 488-labeled anti-Myc antibody added to the culture medium.

**Extended Data Video 2.** Dyl ECM formation and its splitting by cell division (790-800 min, arrows). Myc-Dyl cells were induced and recorded as in Video 1.

**Extended Data Video 3.** Expansion and rupture of Tyn ECM. Myc-Tyn cells were induced by 500 µM CuSO_4_ for 2 days and were imaged with the addition of Alexa 488-labeled anti-Myc antibody. As the cells grew, Tyn ECM was stretched and broken (90 min, arrowheads), and shrunk. Correspond to Extended Data Fig. 2a.

## Supplementary Information

**Supplementary Table 1** Excel file containing the lists of antibodies, plasmids, reagents, experimental models, and the lists of hardware and software.

